# Dissecting the microenvironment around biosynthetic scaffolds in murine skin wound healing

**DOI:** 10.1101/2020.08.10.243659

**Authors:** Chen Hu, Chenyu Chu, Li Liu, Shue Jin, Renli Yang, Shengan Rung, Jidong Li, Yili Qu, Yi Man

## Abstract

Structural properties of biomaterials play critical roles in guiding cell behaviors and influence the immune response against them. We fabricated electrospun membranes with three types of surface topography (Random, Aligned, and Latticed). The aligned membranes showed immunomodulatory ability, and led to faster wound healing, reduced fibrotic response and enhanced regeneration of cutaneous appendages when used in skin wound repair. Based on that, we performed single-cell RNA sequencing analysis on cells from wounded mouse skin in the presence or absence of the Aligned scaffold. Keratinocytes, fibroblasts, and immune cells including neutrophils, monocytes, macrophages, dendritic cells, and T cells showed diverse cellular heterogeneity. More hair follicle progenitor cells, inner root sheath cells (anagen-related) and fibroblast subsets were found in the Aligned group, which corresponded to the improved regeneration of hair follicles and faster wound closure in the presence of scaffold. Immune responses towards the biomaterial differed from that of control group. In aligned samples, infiltrated macrophages and neutrophils were reduced, whereas more effector T cells were recruited. The time course of immune response was possibly advanced towards an adaptive immunity-dominant stage by the scaffold. The microenvironment around scaffold involved intricate interplay of immune cells and cutaneous cells, and wound healing was the comprehensive results of numerous influencing factors working together.

## Introduction

Biomaterials and devices implanted in the body have a broad spectrum of clinical applications including tissue regeneration, cell transplantation, controlled drug release, and continuous monitoring of physiological conditions (1, 2). Among the components of a cell microenvironment, structural features (macroscale, microscale and nanoscale features) play critical roles in guiding cell behaviors (3). Electrospinning technology has been widely applied in preparing scaffolds due to its simplicity, capacity to form fibers on the micro- and nanoscale, structural control of electrospun membranes and cost-effectiveness (4). Electrospun nanofibers could be assembled into well-ordered nanofiber meshes with different morphology, e.g. parallel alignment or latticed patterns of nanofibers (5, 6), and have been preferentially applied to the regeneration of diverse tissues like skin and bone (6, 7). Upon implantation of a material, cells of both the innate and adaptive immune system have a role in the host response (8). Previous studies illustrated the type 1 (pro-inflammatory) immune polarization driven by T helper 1 (Th1) cells, and the induced pro-inflammatory M1 activation of macrophages (stimulated by IL-2 and IFN-γ from Th1 cells). By contrast, in type 2 immune response, T helper 2 (Th2) cells produce cytokines like IL-4 and IL-13, which regulate the polarization of macrophages towards an anti-inflammatory M2 activation (9, 10). More recently, a type 17 immune response was reported to promote chronic fibrosis in tissue around implants (11, 12). In summary, previous studies usually focused on certain types of immune cells, like macrophages, and explored their roles in the host response. However, the immune response is jointly regulated by various immune cells, whose phenotype and function are dictated by external and internal signals. An overview of different immune cells in the microenvironment will aid in comprehensive understanding of the immune responses elicited by scaffolds. Technological advances such as single-cell RNA sequencing (scRNA-seq) (13, 14) have enabled cell populations, functions, and the nuances of their phenotypes in vivo to be studied at a high resolution. By changing the collector, we developed poly(lactic-co-glycolic acid)-fish collagen (PLGA-FC) hybrid electrospun scaffolds with three types of surface topography, i.e. the group with randomly oriented fibers (Random group), the group with mesh-like topography in macroscale and randomly oriented fibers in microscale (Latticed group), and the group with aligned fibers (Aligned group). We explored the regenerative outcomes of these scaffolds in rat/mouse dorsal skin excisional wounds, and evaluated their immunomodulatory properties. The scaffold with the best performance was further investigated. Microenvironment around the scaffold was probed by scRNA-seq. Heterogeneity of keratinocytes, fibroblasts, and immune cell populations, cellular functions, and their interactions in vivo were explored.

## Results

### Evaluation of wound healing in a rat skin wound model

We fabricated scaffolds with random, aligned and latticed patterns of fibers (**Fig. 1A**), and placed them below the full-thickness excisional wound (diameter=6mm) on rat dorsal skin (SD rat). The Ctrl group received no scaffolds (**Fig. 1C**). Biophysical properties of the scaffolds were summarized in **Fig. S1**. Workflow for evaluating wound healing was summarized in Fig. 1B. Wound healing rate was significantly accelerated by the aligned membranes (**Fig. 1D-E**). The Latticed group showed delayed wound healing with the largest residual wound area left at day 7. On day 14, basically all groups achieved complete closure of the wound. Above the scaffolds, surrounding epithelium formed epithelial tongue as the first layer advancing towards the wound (15). On day 7, the Aligned group presented the fastest coverage of the wound, leaving the smallest gap width, whereas the latticed membrane seemed to impede the advancement of surrounding epithelium (**Fig. 1F-G, Fig. S2**). On day 14, re-epithelialization was completed in all groups except for the Latticed group. All groups showed matured stratified epithelium (**Fig. 1F**). Immunofluorescent staining for Krt5 (keratin secreted by keratinocytes in the basal layer) and Krt10 (keratin secreted by differentiated keratinocytes in the supra-basal layers) showed that on day 7, the Aligned group had the largest area of newly formed keratinized epithelium, and more keratinocytes were undifferentiated (Krt5-positive). On day 14, all groups revealed formation of stratified epithelium (**Fig. 1H-I**) (15). For the dermis part, wound space below epithelium was filled with granulation tissue on day 7. Collagen kept being deposited and remodeled (**Fig. S2**). On day 28, the Aligned group had more regenerated hair follicles and sebaceous glands than other treatment groups (**Fig. 1F**). The scaffolds were placed subcutaneously in rats to evaluate host response against them. Thickness of fibrotic capsules around scaffolds was measured at day 3, 7 and 14. The Aligned group had the smallest fibrotic capsule thickness at all time points (**Fig. 1J-K, Fig. S3**). The bulk-tissue RNA-Seq analysis for samples harvested on day 7 (n=3 for each group) identified transcripts corresponding to 34459 genes, distributed over 6 orders of magnitude of expression level. Principal component analysis (PCA) of the data revealed that gene expression profiles of the Random and Aligned samples were more similar, compared to that of Latticed and Ctrl samples (**Fig. 1L**). In Gene ontology (GO) analysis, random and aligned scaffolds induced up-regulated expression of genes associated with immune responses when compared with the Ctrl samples, whereas the Latticed group did not (**Fig. 1M**). Kyoto Encyclopedia of Genes and Genomes (KEGG) analysis also revealed gene enrichment in an immune-related pathway “Cytokine-cytokine receptor interaction” (KEGGID rno04060) (**Fig. S4A-B**) (16). Real-time Quantitative Polymerase Chain Reaction (qPCR) confirmed these elevated genes observed in Aligned and Random groups (**Fig. S4C**). The data suggested that random and aligned membranes were able to modulate the local immune microenvironment. To explore whether the scaffolds had similar performance in other species, we applied them in mouse skin wound model.

**Fig. 1.**
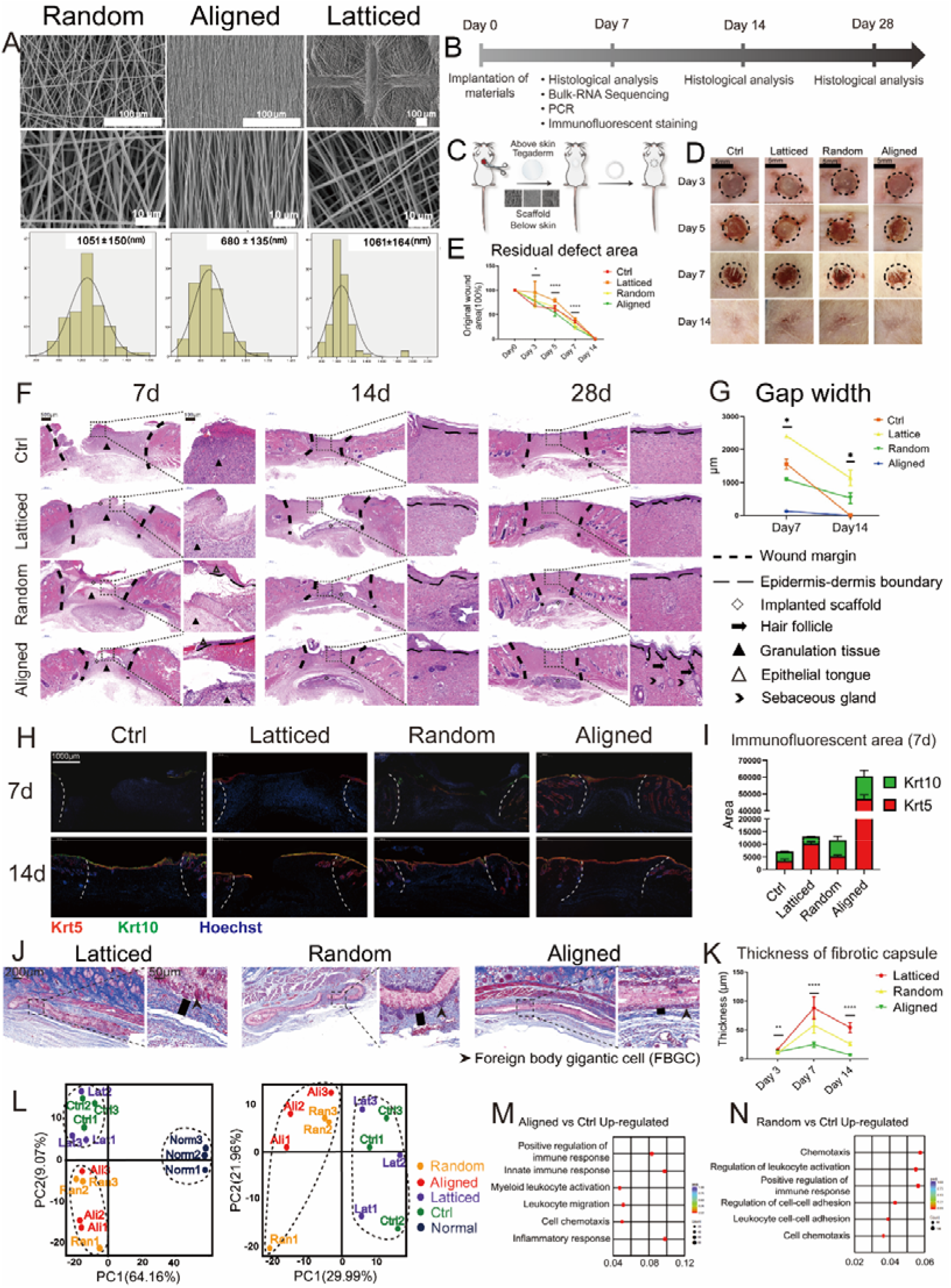
Evaluation of rat skin wound healing implanted with three types of electrospun scaffolds at 7 days post-wounding. (A) Surface topography of three types of electrospun membranes. (B) Workflow for evaluating rat skin wound healing. (C) Surgical processes for the rat skin excisional wound model. (D, E) Residual wound area at each time point. (F) Histological view of Ctrl group and groups implanted with three types of scaffolds. (G) Semi-quantitative evaluation of gap width. (H) Immunofluorescent staining using Krt5 (red) and Krt10 (green). (I) Semi-quantitative evaluation of the fluorescent area. (J) Evaluation of foreign body reaction around biomaterials. FBGCs lined up on Random and Latticed membranes, whereas fewer of them appeared on Aligned ones. (K) Thickness of fibrotic capsules. (L-N) PCA and GO analysis revealed immunomodulatory effects for aligned and random scaffolds. ****P < 0.0001, **P < 0.01, and *P < 0.05 by ANOVA for data in (E), (G) and (K).

### Wound healing in mouse

Workflow for evaluating wound healing in C57BL/6 mouse was summarized in Fig. 2A. We placed three types of scaffolds below the full-thickness excisional wound (diameter=6mm) (**Fig. 2B**). The Ctrl group received no scaffolds. Wound coverage was faster in the Aligned (residual wound area=29.29±4.81%) and Ctrl group (20.80±4.66%) on day 7. On day 14, basically all groups achieved complete closure of the wound (**Fig. 2C-D**). The Aligned group had the smallest gap width on day 7, followed by the Ctrl group (**Fig. 2E-F**). On day 14, re-epithelialization was completed in all groups except for some samples of the Latticed group. All groups showed matured stratified epithelium. Regeneration of hair follicles was seen in Aligned samples after 14 days of healing (**Fig. 2E**). Scaffolds were also placed subcutaneously to evaluate the host response against them. The aligned membranes had the smallest capsule thickness over the observation period (**Fig. 2G, Fig. S5**). Bulk-tissue RNA-Seq for Aligned and Ctrl samples harvested on day 7 (n=3 for each group) was performed. Compared with Ctrl, the Aligned group showed increased genes enriched in inflammatory response, leukocyte chemotaxis and migration (**Fig. 2I**). KEGG analysis revealed elevated gene expression in immuno-related signaling pathways for the Aligned group (KEGGID mmu04657, mmu04668, mmu04060 and mmu04064) (**Fig. 2J**). To summarize, in both rat and mouse models, the aligned membranes led to faster wound healing, reduced fibrotic response and enhanced regeneration of cutaneous appendages compared to other membranes. Meanwhile, they showed immunomodulatory properties. We therefore sought to explore how aligned membranes regulated peri-implant microenvironment. Here, we used scRNA-seq to sequence cells from the wounded murine full-thickness skin 7 days post-wounding.

**Fig. 2.**
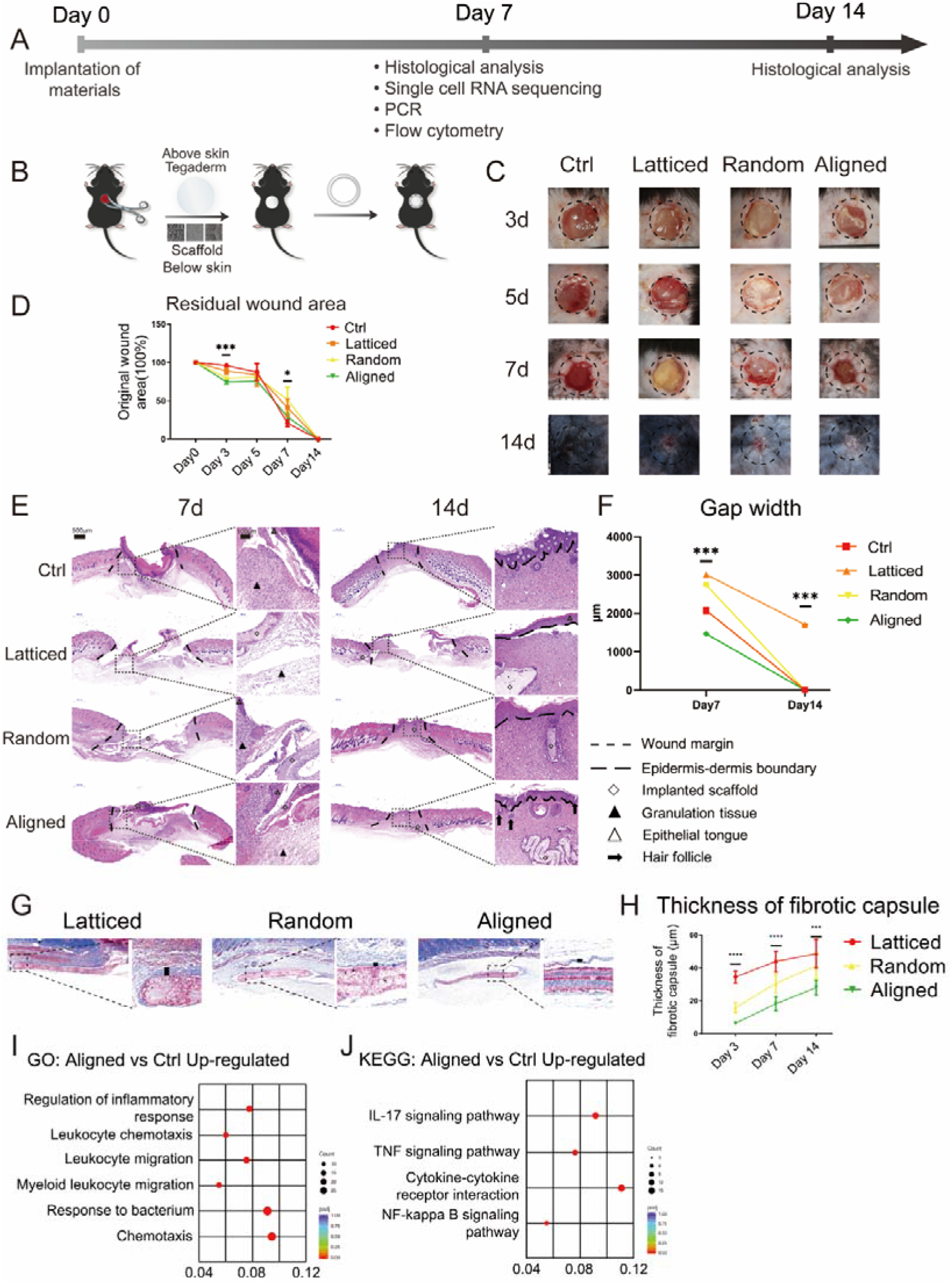
Evaluation of mouse skin wound healing implanted with three types of electrospun scaffolds at 7 days post-wounding. (A) Workflow for evaluating mouse skin wound healing. (B) Surgical processes for mouse skin excisional wound model. (C, D) Residual wound area at each time point. (E) Histological analysis of Ctrl group and groups implanted with three types of scaffolds. (F) Semi-quantitative evaluation of gap width. (G) Evaluation of foreign body reaction around biomaterials. (H) Thickness of fibrotic capsules. (I, J) GO and KEGG analysis revealed immunomodulatory effects for aligned scaffolds. ****P < 0.0001, ***P < 0.001, and *P < 0.05 by ANOVA for data in (D), (F) and (H).

### Single-cell transcriptome analysis of full-thickness skin after wounding and scaffold placement

We isolated cells from the Aligned and Ctrl samples (n=4 biological replicates in each group), and applied them to the 10X scRNA-seq platform (**Fig. 3A**). A total of 8,982 cells in the Ctrl group and 9,593 cells in the Aligned group were captured. After cell filtering, 17,181 single cell transcriptomes were included in the final dataset (8,869 for the Aligned group and 8,312 for the Ctrl group) (**Fig. S6**). We first computationally pooled cells from Ctrl and Aligned groups to create a virtual aggregate. Unsupervised clustering using Seurat categorized the cells into 26 clusters based on global gene expression patterns (Fig. S7), which were assigned to thirteen main classes of cells (**Fig. 3B**): keratinocytes (KER), fibroblasts (FIB), sebocytes (SEB), smooth muscle cells (SMC), endothelial cells (EC), Schwann cells (SC), melanocytes (MEL), innate lymphoid cells (ILC), monocyte-macrophages (MAC), T cells (TC), neutrophils (NEU), dendritic cells (DC) and B cells (BC). Marker genes for each cell cluster were shown in the heatmap (**Fig. 3C**). The composition of each cell cluster was listed so that the proportion of cells from two groups could be identified across all cell clusters (**Fig. 3B**). As smooth muscle cells (57% from the Aligned group and 43% from the Ctrl group) could hardly regenerate at 7 days post-wounding, we regarded a proportion within 50±7% as equilibrium between two groups. Genes related to macrophages (Itgam, Cd68, Arg1, Mrc1) were significantly higher expressed by the Ctrl samples (**Fig. 3D**).

**Fig. 3.**
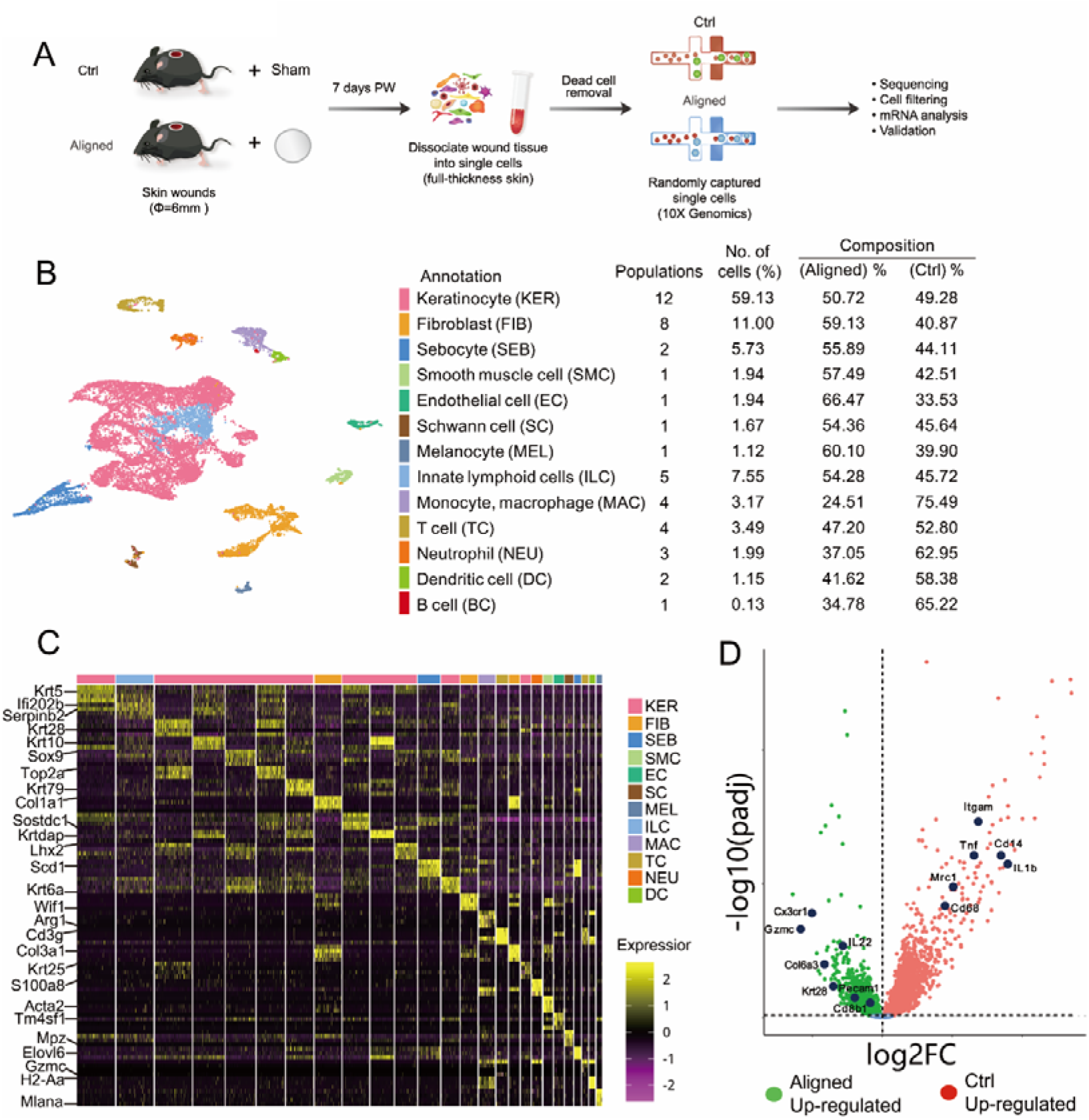
Overview of the single-cell transcriptome analysis. **(A) Workflow for single-cell experiment**. (B) Cells were categorized into thirteen main classes. Cell populations in each class, number of cells (%), and composition of aligned and Ctrl cells were listed. (C) Marker genes for different cell classes. (D) Gene expression differences between Ctrl and Aligned groups.

### Sub-clustering of keratinocytes reveals a higher proportion of hair follicle progenitor cells from the Aligned group

We selected cells that were in the first-level clustering defined as keratinocyte, and subjected them to a second round of unsupervised clustering (**Fig. 4A**). Terminal cornification of epidermis is achieved by keratinocytes passing through basal layers, differentiated layers, and cornified layers (15). In our study, the inter-follicular epithelium was composed of the basal layer cells (IFEB, Krt5^hi^Krt14^hi^), supra-basal layer cells (Krt10^hi^Krt1^hi^ IFED1, Krt10^hi^Mt4^hi^ IFED2), and cornified layer (Lor^+^ IFED3) cells (**Fig. 4A-B**). The location of keratinocyte subsets was marked in **Fig. 4C**. Hair follicles in this study were in second telogen since the mice were 8 to 10 weeks old when they were sacrificed (17). However, anagen hair follicle gene signatures were also found, possibly resulting from hair follicle regeneration after wounding. Upper hair follicle cells were separated into three subsets named uHF1 (Krt79^hi^Krt17^hi^), uHF2 (Klk10^+^Krt79^hi^), and uHFB (the upper hair follicle basal layer cells, Sostdc1^hi^Apoe^hi^). The hair follicle progenitor cells (Krt28^hi^Lhx2^hi^Mki67^hi^ HFP) were highly proliferative. Germinative layer cells (Mt2^hi^Dcn^hi^) belonging to anagen hair follicles also expressed high cell proliferation related genes like Top2a and Birc5. Meanwhile, they had a basal cell gene signature (Krt5^hi^Krt14^hi^). The inner root sheath (IRS) and cortex cells were characterized by Krt28, Krt27, Krt73, and Krt25, markers for the Henle and Huxley layers of anagen hair follicles (18). Cells of the inner bulge layer (IB) and outer bulge layer (OB) expressed their typical gene signatures (**Fig. 4A-C**). When analyzing the inter-group differences, the Aligned group contributed to a larger proportion of hair follicle progenitor cells (HFP) and highly proliferative inner root sheath cells (IRS) (**Fig. 4A**). The overall difference analysis also revealed up-regulation of genes (Krt28, Sox18) related to hair follicle stem cells in the Aligned group (**Fig. 3D**). This might explain the enhanced hair follicle regeneration observed in Aligned groups either on rat or mouse models.

**Fig. 4.**
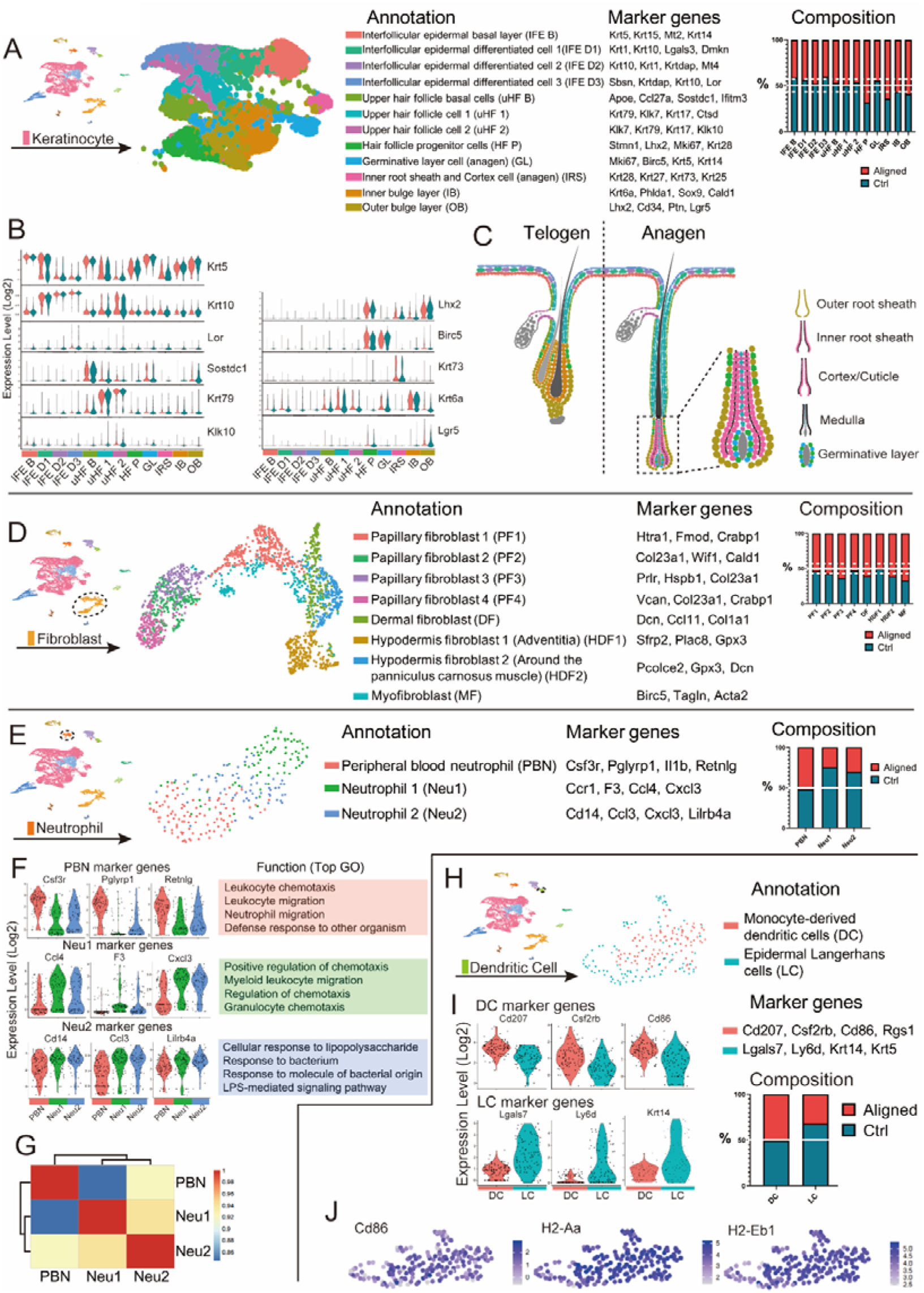
Further analysis of keratinocytes, fibroblasts, neutrophils and dendritic cells. (A) Sub-clustering of keratinocytes revealed twelve subsets. Marker genes and composition for each subset was listed. (B) Marker genes for each keratinocyte subset. (C) Location of different keratinocyte subsets. Color of each cell corresponds with its cell type in (A). (D) Sub-clustering of fibroblasts revealed eight subsets. The marker genes and composition for each subset was listed. (E) Sub-clustering of neutrophils revealed three subsets. (F) Marker genes for neutrophil subsets and their enriched gene sets in GO analysis. (G) Correlation analysis of neutrophil subsets. (H) Sub-clustering of dendritic cells revealed two subsets. (I) Marker genes for dendritic cell subsets. (J) Expression of genes associated with antigen presenting in dendritic cells.

### Inter-group differences in fibroblasts suggest more active ECM formation in the Aligned group

The dermis consists of several layers: the papillary dermis lies closest to the epidermis, the underlying reticular dermis contains the bulk of the fibrillary extracellular matrix, and beneath the reticular dermis lies the hypodermis (19). Fibroblasts from different layers presented distinct gene signatures (**Fig. 4D**). There were four populations of papillary fibroblasts (Crabp1^+^Col23a1^+^) (20, 21). Dermal fibroblasts expressed increased Ccl11 and Dcn. The Gpx3^+^Plac8^hi^ subset was identified as hypodermis fibroblast located to the adventitia (HDF1), and the Gpx3^+^Plac8^lo^ subset (HDF2) was around the panniculus carnosus muscle (18). Contractile myo-fibroblasts (Acta2^+^ MF) expressed elevated genes associated with cell proliferation (Birc5^hi^). Composition of each subset revealed that more papillary fibroblasts (PF2), dermal fibroblasts (DF), hypodermis fibroblasts (HDF2) and myo-fibroblasts (MF) were from the Aligned group, suggesting more robust ECM formation in the presence of scaffolds.

### Neutrophils and dendritic cells are more abundant in the Ctrl samples

After wounding, neutrophils close to the focus of injury migrate toward the nidus, followed by those recruited from more than 200μm from the site of tissue injury (22). Three neutrophil subsets were identified in our study (**Fig. 4E**). Peripheral blood neutrophils (PBN) expressed typical gene signatures including Csf3r, Pglyrp1, Il1b, and Retnlg. Neutrophil 2 (Neu2) expressed elevated Cd14 and Ccl3, and was identified as antimicrobial phagocytic neutrophils. Neutrophil 1 (Neu1) expressed elevated Ccr1, a chemokine receptor that mediates neutrophil migration. Correspondingly, PBN showed gene enrichment in leukocyte chemotaxis and migration; Neu1 expressed genes enriched in leukocyte/granulocyte chemotaxis and migration, and Neu2 was enriched in anti-bacterial biological processes (**Fig. 4F**). According to the correlation analysis, Neu 1 and Neu 2 were derived from PBN in circulation (**Fig. 4G**). In Neu1 and Neu2, more cells belonged to the Ctrl group (**Fig. 4E**), suggesting more significant Neu1 and Neu2 infiltration in the Ctrl samples at a proliferative stage (7 days post-wounding). Dendritic cells were classified into two subpopulations (**Fig. 4H**). The subset derived from monocytes (DC) was characterized by increased Cd207 and Cd86 (10, 23). The Langerhans cell subset bore a keratinocyte gene signature, probably transferred from the resident microenvironment (**Fig. 4I**) (24). Both DC and LC presented elevated major histocompatibility complex (MHC) molecules, suggesting an antigen presenting function of them (**Fig. 4J**). LCs contained more Ctrl-derived cells, indicating differences in Langerhans cell infiltration between groups.

### Macrophage heterogeneity and their down-regulation by scaffolds

To explore the heterogeneity of macrophages in vivo and their inter-group differences, we subjected them to further unsupervised sub-clustering. Four subsets were determined (**Fig. 5A**). The subset that showed increased anti-inflammatory genes (Ccl8, Folr2, C1qa and Mrc1) was named anti-inflammatory macrophages (AIM) (25, 26). Another subset characterized by elevated expression of pro-inflammatory genes (Ptgs2, Ccl3, Inhba, Nos2) was named pro-inflammatory macrophages (PIM) (27, 28). The monocyte subset (Mono) showed higher Ly6c2, Plac8, Cd14, and Clec4e, indicting inflammatory responses against lesions and microorganism (29, 30). The Cytip^hi^H2-Eb1^hi^ subset of cells was defined as monocyte-derived dendritic cells (M-DC) (**Fig. 5B**) (31). We further found that canonical M1 and M2 markers were not entirely consistent with computationally determined AIM and PIM. Arg1, a canonical M2 marker, was expressed by basically all monocyte-macrophage subsets (AIM, PIM, Mono), and was regarded as a pan-macrophage marker in this study (**Fig. 5B**). Expression of another type 2 gene, Socs3, did not parallel Mrc1 expression either. Similar disparity was found in the expression of canonical type 1 genes. M-DC, rather than PIM, expressed more Cd86. Neither was the expression of Cd86 correlated with Nfkbiz. Expression of other genes associated with fibrotic or regenerative macrophage subsets in a scaffold immune microenvironment did not correspond with these clusters either (**Fig. 5B**) (12). It was determined that Ly6c2, Arg1, Mrc1 and Nos2 were sufficient to distinguish the computationally determined AIM, PIM and Mono subsets. We performed flow cytometry on cells isolated from the Aligned and Ctrl samples using Cd68 (a monocyte-macrophage marker, also expressed by some neutrophils and dendritic cells) and the proposed markers (Ly6c2, Arg1, Mrc1 and Nos2). The Cd68^+^ cells were selected to create a t-distributed stochastic neighbor embedding (tSNE) plot. We then identified Arg1^+^ macrophages expressing the surface markers Mrc1 and Nos2 in the gated dataset to represent AIM and PIM, respectively. The Arg1^+^Ly6c2^+^ monocytes were also identified (Mono). The three terminal clusters (AIM, PIM, and Mono) could be separated, indicating that the subsets can be identified experimentally using flow cytometry (**Fig. 5C**). Pseudo-temporal trajectory (Monocle 2) of the four subsets revealed that monocytes developed into M-DCs and polarized macrophages (AIM and PIM). Although AIM and PIM expressed distinct gene signatures, they were highly correlated (**Fig. 5D-E**). The Ctrl samples contributed to a larger proportion of cells in all subsets (**Fig. 5A**). Therefore, the scaffold might play a role in the reduction of macrophage infiltration at the proliferative stage.

**Fig. 5.**
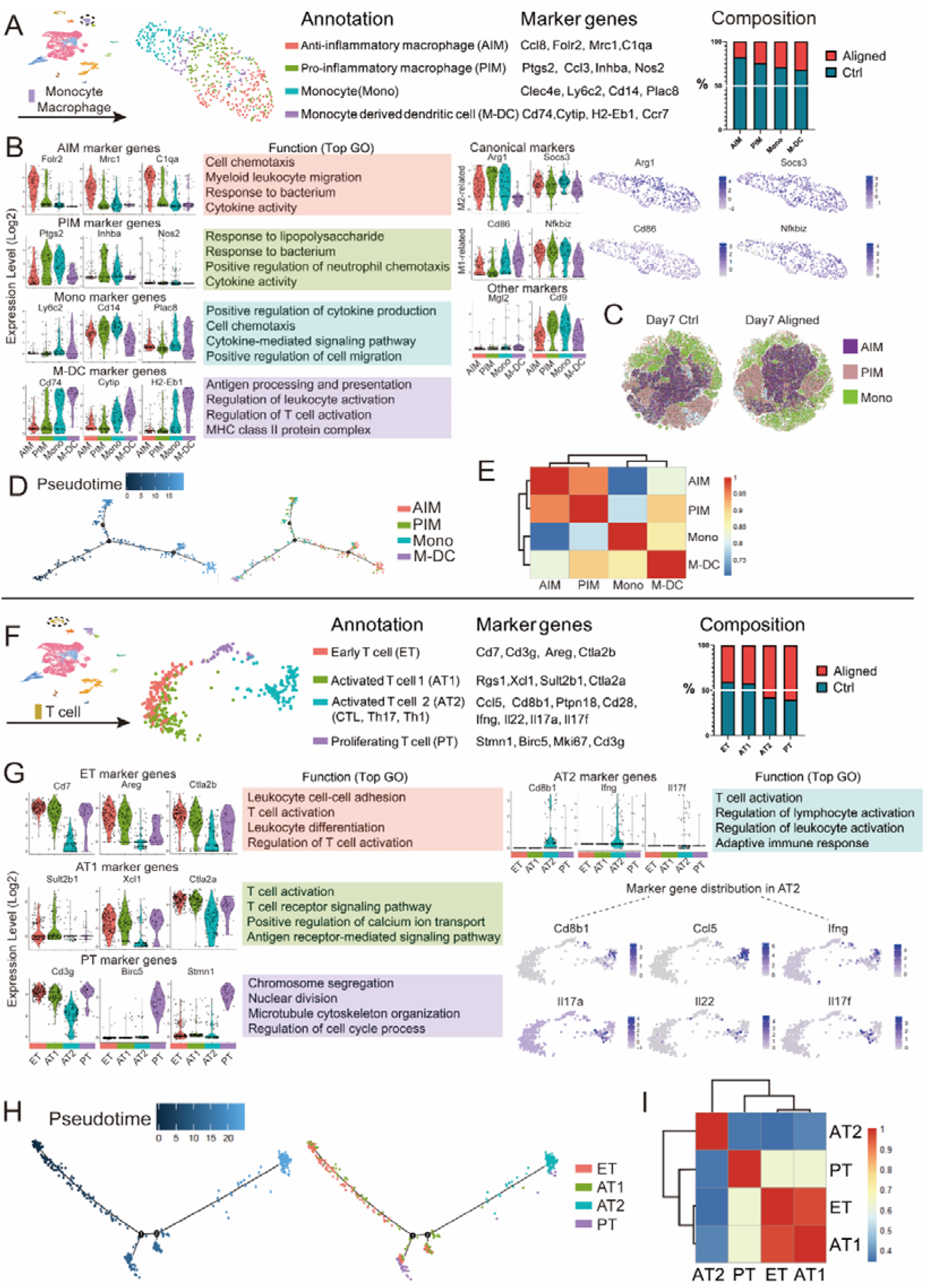
Further analysis of macrophages and T cells. (A) Sub-clustering of macrophages revealed four subsets. Marker genes and composition for each subset was listed. (B) Marker genes for each macrophage/monocyte subset and their enriched gene sets in GO analysis. Expression of canonical M1 and M2 markers and other proposed markers were shown. (C) In vivo flow cytometry strategy using Cd68 to identify monocytes and macrophages. Arg1+ macrophages expressing the surface markers Mrc1 and Nos2 in the gated dataset represented AIM and PIM, respectively. The Arg1+Ly6c2+ monocytes were also identified (Mono). (D) Pseudo-temporal ordering of macrophage/monocytes, and the distribution of four subsets along the trajectory. (E) Correlation analysis of macrophage/monocyte subsets. (F) Sub-clustering of T cells revealed four subsets. (G) Marker genes for T cell subsets and their enriched gene sets in GO analysis. AT2 subset expressed gene signatures of cytotoxic T cells, Th1 cells, and Th17 cells. (H) Pseudo-temporal ordering of T cells, and the distribution of four subsets along the trajectory. (I) Correlation analysis of T cell subsets.

### Sub-clustering of T cells revealed a novel T cell population and more effector T cells in the Aligned group

T cells were clustered into four subsets (**Fig. 5F**). T cells characterized by increased Cd7, Cd3g, and Areg were named as Early T cells (ET) (32). The subset adjacent to ET expressed elevated Xcl1 and Sult2b1 (associated with T cell activation), and was named Activated T cell 1 (AT1) (33). AT1 also expressed increased Areg, Ctla2a, and Ctla2b, genes related to immune homeostasis and immuno-suppression (34, 35). Another activated T cell subset (AT2) expressed up-regulated genes associated with cytotoxic T cells (Cd8b1), Th1 cells (Ifng, Ptpn18), and Th17 cells (Il17a, Il17f) (36, 37). Therefore, AT2 might include multiple effector T cells. A novel T cell subset connecting AT2 and ET was featured by high expression of Birc5, Mki67 and Stmn1, markers for cell proliferation, and was named Proliferating T cell (PT) (**Fig. 5F-G**). In GO analysis, both ET and AT1 were enriched in T cell activation. AT1 was also found elevation of T cell receptor signaling pathway, suggesting that these T cells played a role in antigen recognition (**Fig. 5G**). We found three terminally differentiated clusters stemming from two precursors (ET and AT1) in pseudo-time analysis. PT was differentiated from AT1 and ET after the first branch point, and might be a transitional status between early T cells and effector T cells. At the second branch, cells differentiated into two terminal clusters. One belonged to AT1 and ET, and another one was the AT2 population (**Fig. 5H**). Correlation analysis showed that ET and AT1 were highly correlated (**Fig. 5I**). In the ET and AT1 populations, more cells were from Ctrl samples, whereas a larger number of cells in AT2 and PT were from aligned samples.

### Receptor-ligand analysis reveals intricate interactions among immune cells, keratinocytes and fibroblasts

To explore potential interactions among immune cells, keratinocytes and fibroblasts, we ran CellChat analysis on these datasets (38). For anti-inflammatory macrophages (AIM), broadly speaking, “pro-inflammatory” signals like Tnf, Visfatin, Rankl, EGFR and C3a signaling, and “anti-inflammatory” signals including Spp1, Lgals9, Sema3, Chemerin, Il2, Il13 and Il10 signaling were involved either in a paracrine or autocrine way. Signals like Mif, Ccl, Csf, Edn, and Cxcl12 signaling exerted protective or deleterious effects depending on their specific roles (**Fig. 6E**) (25, 39–50). Among T cell populations (**Fig. 6F**), ET communicated with other T cells mainly through Tgfb signaling, mediating ET cell tolerance and immune homeostasis (51). The novel PT population secreted Wnt11 signals that bound to Fzd5 receptor on ET. Inner bulge layer keratinocytes secreted Cxcl16 that targeted Cxcr6 on ET, chemoattracting ET to the wound area (52). For fibroblast populations, HDF1 had the most frequent interaction with other cells (**Fig. 6G**). In addition to the mentioned signals, OSM signaling was also activated and induced the expression of the pro-inflammatory cytokine IL-6 (53). T cells and macrophages secreted Tgfb1 and Tgfb2 to target HDF1 and PF1, leading to proliferation and differentiation of fibroblasts into myo-fibroblasts (54). Among keratinocytes, IFEB and IB1 had the most frequent contact with other cells (**Fig. 6H, Fig. S8I-M**). IFEB received Tgfb1 and Tgfb2 (signals playing key roles in regulating epithelial-to-mesenchymal transition (EMT) for keratinocytes) from T cells, macrophages and germinative layer (GL) keratinocytes (55). Interactive plots for other types of cells were summarized in Fig. S8. The communication between immune cells and cutaneous cells was complicated, and wound healing was the comprehensive results of numerous influencing factors working together.

**Fig. 6.**
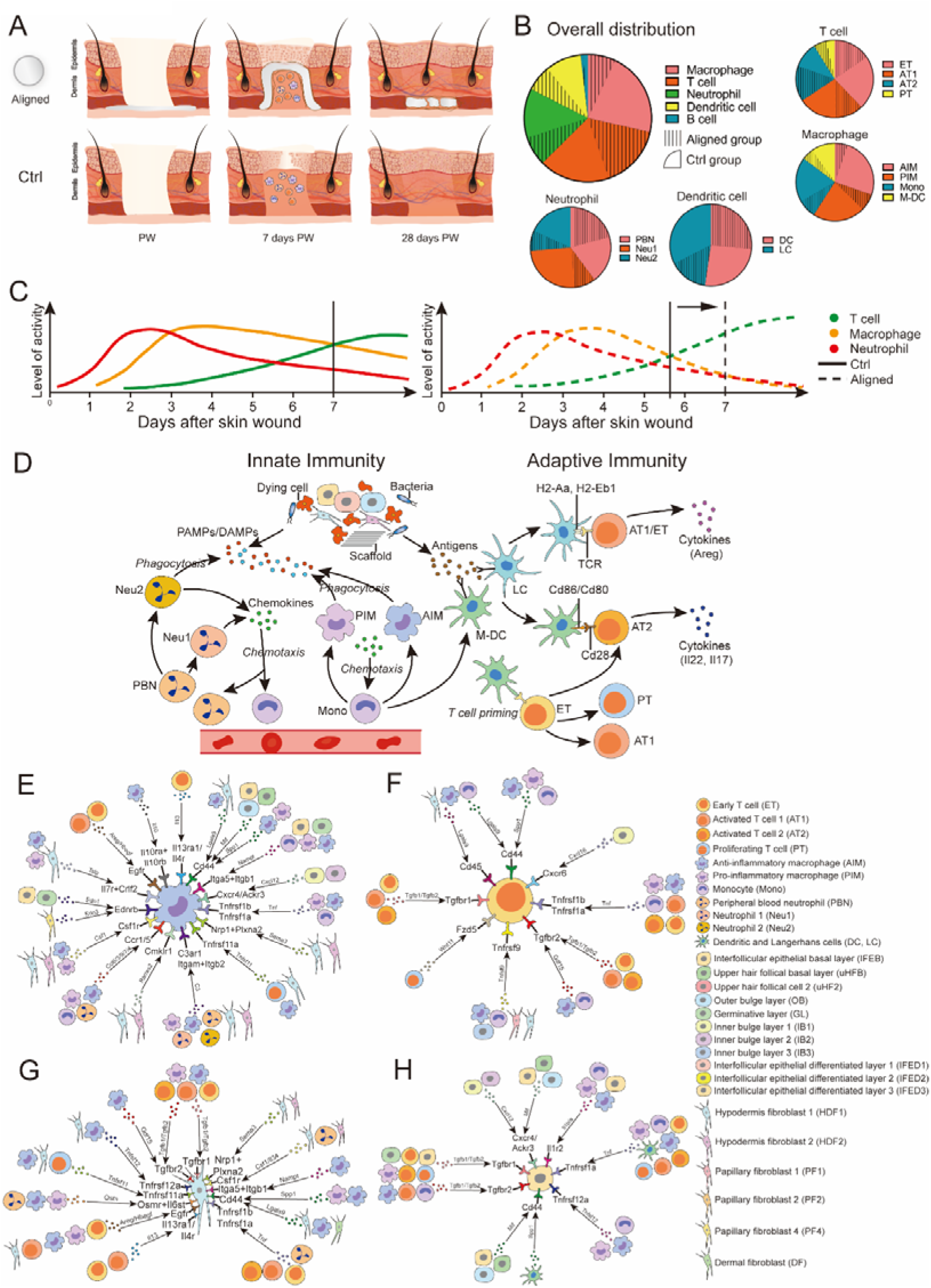
Summary of the differences in wound healing and immune responses between the Aligned group and Ctrl group. (A) The aligned membranes had immunomodulatory properties, and led to faster wound healing, reduced fibrotic response and enhanced regeneration of skin appendages. (B) Pie plot for overall immune cells in two groups. For each cell type, differences in cell amount between aligned and Ctrl groups. (C) Timeframe of innate and adaptive immune responses occurred in Aligned and Ctrl groups. (D) The immune microenvironment around aligned scaffolds. (E) Cell-cell interaction plots according to receptor-ligand analysis via CellChat.

## Discussion

For this study, we fabricated electrospun membranes with three types of surface topography (Random, Aligned, and Latticed). The aligned membranes led to faster wound healing, reduced fibrotic response and enhanced regeneration of cutaneous appendages compared to other scaffolds. Meanwhile, the aligned membranes exhibited immunomodulatory properties (**Fig. 6A**). Based on that, we generated single-cell transcriptomes from wounded mouse skin to investigate the microenvironment around scaffolds. The skin wound repair process was classically divided into four phases: hemostasis (hours), inflammation (days), proliferation (1-2 weeks), and remodeling (>2 weeks). 7 days post-wounding was a transitional time point when the innate immune response subsided and activity of adaptive immune cells increased (10). In the Ctrl samples, the infiltration of T cells achieved a similar extent with macrophages (**Fig. 6B** Overall distribution). However, in Aligned group, infiltrated macrophages were much fewer than T cells, and more terminally differentiated effector T cells were present (**Fig. 6B**). According to the timeframe of innate and adaptive immune responses (10), the process of innate immunity seemed to be alleviated earlier, and adaptive immune response was advanced in the presence of aligned scaffolds (**Fig. 6C**). In the immune microenvironment around aligned scaffolds (**Fig. 6D**), damage-associated molecular patterns (DAMPs), pathogen-associated molecular patterns (PAMPs) and antigens from cell debris, pathogens, and foreign agents (scaffold) triggered innate and adaptive immune responses. Neutrophils from circulation (PBN) quickly migrated to the wound area. The infiltrated neutrophils (Neu1 and Neu2) phagocytosed dying cells and microorganisms, and secreted chemo-attractants like Ccr1 and Ccl4 to recruit more leukocytes and lymphoid cells (52). Circulating monocytes (Mono) were also recruited to the wound area, and differentiated into pro-inflammatory macrophages (PIM), clearing dying neutrophils and debris. Meanwhile, anti-inflammatory macrophages (AIM) chemo-attracted more cells via chemokines like Ccl8 and Ccl6. Part of monocytes differentiated into dendritic cells (M-DC). Tissue resident Langerhans cells (LC), together with M-DC, functioned as antigen-presenting cells (APCs). In the antigen specific signal, LC and DC processed antigens into peptides, and presented them by MHC class II molecules (H2-Aa, H2-Eb1) on the cell surface. T cells (ET and AT1) bound to the MHC molecules through surface receptors (TCR), and differentiated into effector T cells (Cd4^+^ AT2 cells). The co-stimulatory signal was characterized by the engagement of Cd28 receptor on T cells (AT2, Cd8b1^+^Cd28^hi^) with Cd86 ligands on APCs (Cd86^+^ DC and LC) (10, 56). With regard to the influences that immune microenvironment had on tissue generation, immune cells sent out a variety of signals (like Grn, Tgfb, Areg/Hbegf) that modulated behaviors of keratinocytes and fibroblasts (**Fig. 6**). On the other hand, keratinocytes and fibroblasts secreted chemotactic, pro- or anti-inflammatory signals that regulated immune cell polarization and function. Immune cells themselves also interact with each other via numerous signals. The communication network of immune cells and cutaneous cells was complicated, and wound healing was the comprehensive results of these factors.

## Acknowledgement

This work was supported by Research and Develop Program, West China Hospital of Stomatology Sichuan University (No.LCYJ2019-19); The Fundamental Research Funds for The Central Universities (No. 2082604401239); National Key Research and Development Program of China (No. 2016YFA0201703/2016YFA0201700)

## Materials and methods

### Electrospinning of polymer scaffolds

We used poly(lactic-co-glycolic acid) (PLGA) (LA/GA = 75:25, Mw = 105 kDa, dispersity is 1.897) produced by Jinan Daigang Biomaterial Co., Ltd. (Shandong, China) and FC (from fish scale and skin) obtained from Sangon Biotech Co., Ltd. (Shanghai, China) to fabricate scaffolds by electrospinning. The PLGA (20% w/v) and FC (2% w/v) solution (dissolved in 1,1,1,3,3,3-Hexafluoro-2-propanol (HFIP) solvent (Aladdin Co., Ltd. (Shanghai, China)) were loaded into a plastic syringe fitted with a flat-tipped 21G needle (inner diameter=0.5mm). A high voltage of 7kV and a distance of 16 cm were applied between the needle and the collector. For randomly oriented fibers (Random group), the electrostatically charged fiber was ejected toward the grounded flat collector in the high electric field, forming a membrane deposited on the aluminium foil. For the Latticed group, an electroconductive chess-like wire net was used as the collector. For the Aligned group, the rotational speed of the collecting drum is set at 2800rpm. Finalized scaffolds were approximately 30–90 μm in thickness. To crosslink FC, the membranes were immersed in 50 mM of EDS/NHS and 10 mM of MES ethanol solution for 24 h at 4 °C. Then membranes were washed three times with 75% ethanol and dried in vacuum oven for 24 h. Subsequently, the prepared membranes were sterilized by γ-irradiation for in vitro and in vivo experiments.

### Characterization of scaffolds

Scanning electron microscopy (SEM; JEOL, JSM-6510LV, Japan) was employed to observe the surface morphology of the electrospun membranes. Image-Pro Plus was applied to quantitatively measure the fiber diameter and distribution from the SEM images obtained. The surface wetting behavior of the membranes were characterized by measuring the water contact angles (Chengde Dingsheng, JY-82B, China). Five samples were tested for each type of membrane to obtain an average value. The tensile properties of the membranes were tested under a constant upper clamp at speed of 15 mm/min. All tensile tests follow the criteria of “Plastics-Determination of tensile properies of films” (GB/T 1040.3-2006, corresponding with ISO 1184-1983).

### Cell culture and cell viability test

L929 mouse fibroblast cells and Human oral keratinocytes (HOK) were used for viability tests. Cells were cultured in medium containing RPMI 1640 medium (HyClone) supplemented with 10% fetal bovine serum (Gibco), and were kept at 37□ in humidified 5% CO2/95% air. The cell viability was determined by Cell Counting Kit-8 (CCK-8, Dojindo Laboratories, Kumamoto, Japan). Electrospun membranes were cut into squares (edge length=5mm) and placed in the bottom of 96-well plates (n=3 for each group). L929 cells and HOK cells were seeded onto membranes at 4*10^4^ cells/ml. Cells were co-cultured with membranes for 1, 3, and 5 days. Blank wells seeded with equal amount of cells were used as Ctrl. 10μl CCK-8 solution was added to each well, and the plates were incubated at 37 °C for 1h. After incubation, the Abs at 450nm was measured to determine the cell viability using a micro-plate reader (Multiskan, Thermo, USA).

### Experimental model

#### Excisional wound model

The protocol of the present experiment was approved by Institution Review Board of West China Hospital of Stomatology (No.WCHSIRB-D-2017-033-R1) Animals included Sprague Dawley male rat at ages from 7 to 8 weeks and C57BL/6 male mice at ages from 7 to 9 weeks (Chengdu Dossy Experimental Animals Co., LTD.). Hair at the surgical area was removed. Full-thickness circular excisional wound (diameter=6mm) was created at the dorsal skin of rats/mice. Random, aligned and latticed electrospun scaffolds were trimmed into circular shape (diameter=8mm), and placed below the wound. The Control group did not receive any implants. A sterile Tegaderm film (3M) was placed above the wound to protect the wound area. Then annular silicone splints (inner diameter=8mm, outer diameter=12mm) were sutured with the Tegaderm film and underlying skin in order to minimize the contraction of the dorsal muscle. After healing for 1, 2 and 4 weeks, animals were euthanized for sample harvest. Using the residual wound as center, a round skin sample (diameter =10mm) containing all the layers of skin was harvested.

### Model for subcutaneous implant placement

The surgical area on dorsal skin was shaved and aseptically prepared. Three horizontal incisions of approximately 10 mm were made and subcutaneous pockets were created for membrane implantation. Then random, align and lattice scaffolds were placed into the pockets. After implantation, the incisions were sutured with interrupted sutures. After recovering for 3, 7, and 14 days, samples including scaffolds and the whole layer of skin at surgical sites were together harvested.

### Specimen harvest for scRNA-seq

We obtained skin samples by cutting off skin at the wound area (circular, diameter=10mm). Subcutaneous tissues were removed. A total of four samples were harvested in each group. The tissues were washed in a 100 mm petri dish containing 20 ml of phosphate-buffered saline (PBS). Then they were transferred to a 50 mm petri dish containing 100μL of Enzyme G (Epidermis Dissociation Kit mouse, Miltenyi) and 3.9 ml of PBS buffer with the dermal side facing downwards. Tissues were digested for 16 hours at 4°C. Then they were transferred into a 50 mm petri dish containing 4mL of 1×Buffer S (Miltenyi). Epidermis was peeled off from the skin using tweezers, and was cut into pieces. Enzyme mix containing 3.9 ml of 1×Buffer S, 100 μl of Enzyme P, and 20 μl of Enzyme A (Miltenyi) stored in a gentleMACS™ C Tube was used to digest the epidermis pieces for 20 minutes at 37□. Then 4 ml of PBS that contained 0.5% bovine serum albumin (BSA) was added. A gentleMACS Dissociator (Miltenyi) was applied to automatically dissociate the epidermis (Program B). The sample was passed through a 70μm cell strainer (Corning), centrifuged at 300×g for 10 minutes at room temperature, and resuspended with PBS that contained 0.5% BSA. Cells were gently washed twice and stored in an ice box. For the dermis part, they were first cut into pieces (diameter < 1mm). The tissue was mixed with 10ml enzyme mix containing Type I Collagenase (3125u/mL) (Gibco) and 2.5ml trypsin (Gibco), and poured into a gentleMACS™ C Tube. After dissociating the tissue on gentleMACS Dissociator for 37s (Skin mode), another 10ml enzyme mix was added. The sample was digested for 2.5 hours at 37□ in a rotary machine (Peqlab). Then the dermis sample was passed through a 70μm cell strainer (Corning), centrifuged at 300×g for 5 minutes at room temperature, and resuspended with 3ml red blood cell lysis buffer (Solarbio). After 3 minutes, the cell suspension was centrifuged and gently resuspended with RPMI 1640 medium (Hyclone). Cells were gently washed twice with PBS containing 0.5% BSA and stored in an ice box. The epidermis and dermis cell solutions were mixed together as a whole. The sample was centrifuged, and resuspended with 100 μl Dead Cell Removal MicroBeads (Miltenyi). After incubation for 15min at room temperature, the cell suspension was diluted in 3ml 1×Binding buffer (Miltenyi). LS columns (Miltenyi) and a magnetic stand (Miltenyi) were used for removal of dead cells and debris. The negatively selected live cells passed through the column, and were resuspended with PBS containing 0.05% BSA. Finally, we proceeded with the 10x Genomics^®^ Single Cell Protocol.

### Single-cell encapsulation and library generation

Single cells were encapsulated in water-in-oil emulsion along with gel beads coated with unique molecular barcodes using the 10× Genomics Chromium Single-Cell Platform. For single-cell RNA library generation, the manufacturers’ protocol was performed. (10×Single Cell 3’ v3) Sequencing was performed using Illumina 1.9 mode with 94574 reads per cell.

### Quality control

We use fastp to analyze basic statistics on the quality of the raw reads.

If clean reads is indispensable, then raw read sequences produced by the Illumina pipeline in FASTQ format were pre-processed through Trimmomatic software which can be summarized as below:

1. Remove low-quality reads: scan the reads with a 4-base wide sliding window, cut when the average quality per base drops below 10 (SLIDINGWINDOW: 4:10)
2. Remove trailing low quality or N bases (below quality 3) (TRAILING: 3)
3. Remove adapters: there are two modes to remove the adapter sequence: a. alignment with the adapter sequence, the number of matching bases was greater than 7 and mismatch=2; b. when read1 and read2 overlapping base scoring was greater than 30, we removed non-overlapping portions (ILLUMINACLIP:adapter.fa:2:30:7)
4. Drop reads below the 26 bases long
5. Discard those reads that did not form pairs.

The remaining reads that passed all the filtering steps was counted as clean reads and all subsequent analyses were based on this.

#### Generation and Analysis of Single-Cell Transcriptomes

Raw reads were demultiplexed and mapped to the reference genome by 10X Genomics Cell Ranger pipeline (https://support.10xgenomics.com/single-cell-gene-expression/software/pipelines/latest/what-is-cell-ranger) using default parameters. All downstream single-cell analyses were performed using Seurat. In brief, for each gene and each cell barcode (filtered by CellRanger3.0), unique molecule identifiers were counted to construct digital expression matrices. Secondary filtration by Seurat: A gene with expression in more than 3 cells was considered as expressed, and each cell was required to have at least 200 expressed genes. We also filtered out some of the foreign cells.

The Seurat package was used in data normalization, dimension reduction, clustering, and differential expression. We used the Seurat alignment method canonical correlation analysis (CCA) [Nat. Biotechnol. 36, 411–420 (2018).] for integrated analysis of datasets. For clustering, highly variable genes were selected and the principal components based on those genes used to build a graph, which was segmented with a resolution of 0.6.

### RNA-seq analysis

The bulk tissue RNA-seq analysis was carried out by Novogene Corporation (Beijing, China). RNA was extracted from tissues or cells using standard methods to make sure samples were strictly controlled for quality. In terms of library construction and quality control, mRNA can be obtained in two main ways: firstly, most eukaryotes’ mRNA has poly A-tailed structural, and poly A-tailed mRNA can be enriched by Oligo (dT) magnetic beads. The other is the removal of ribosomal RNA from the total RNA to obtain mRNA. Subsequently, the obtained mRNA was randomly interrupted by divalent cations in NEB Fragmentation Buffer, and the database was constructed according to the NEB general database construction method or chain specific database construction method. Upon completion of library construction, a Qubit2.0 Fluorometer was used for initial quantification, and the library was diluted to 1.5ng/ul. Then the insert size of the library was detected using Agilent 2100 bioanalyzer. After the insert size met the expectation, the effective concentration of the library was accurately quantified by qRT-PCR (the effective concentration of the library higher than 2nM) to ensure library quality. Finally, the libraries were qualified for sequencing, and Illumina sequencing was performed after pooling the different libraries according to the requirements of effective concentration and target data volume, of which the basic principle is Sequencing by Synthesis. Through z-transformation of Fragments Per Kilobase of transcript per Million mapped reads (fpkm) of the selected gene, gene expression was analyzed. Sample size for conventional, bulk RNA-Seq libraries was fixed at 3 biological replicates.

### Quantitative Real-Time Polymerase Chain Reaction (qPCR)

The harvested samples were cut into pieces, and homogenated in TRIzol™ Reagent (Cat. #15596026, Invitrogen, Thermo Scientific). Concentration and ratio of total RNA were detected by NanoPhotometer NP80 (Implen, Westlake Village, CA) at wavelength of 260 nm and 280 nm. The cDNAs were synthesized using PrimeScript™ RT reagent Kit with gDNA Eraser (Perfect Real Time) (Cat. #RR047A), then amplified by qPCR with the specific primers (Table 1). PCR was performed on QuantStudio 3 Real-Time PCR Systems (ThermoFisher Scientific, Waltham, MA). Each 20 μL of PCR mixture contained 10μl of TB Green Premix Ex Taq (Tli RNaseH Plus, Takara) (2X), 0.4μL of PCR Forward Primer (10μM), 0.4μl of PCR Reverse Primer (10μM), 0.4μl of ROX Reference Dye (50X), 2μl Template and 6.8μl of Sterile purified water. Samples were incubated at 1 cycle of 95□ for 30 s followed by 40 cycles of 95□ for 5 s and 60□ for 34 s, and ended up with a cycle composing of 95□ for 15 s, 60□ for 1 min and 95□ for 15 s. Results were analyzed using the comparative CT (ΔΔCT) method to calculate gene expression fold changes normalized to the levels of Gapdh gene transcripts. The experiments were repeated for three times independently (n=3).

**Table1.**
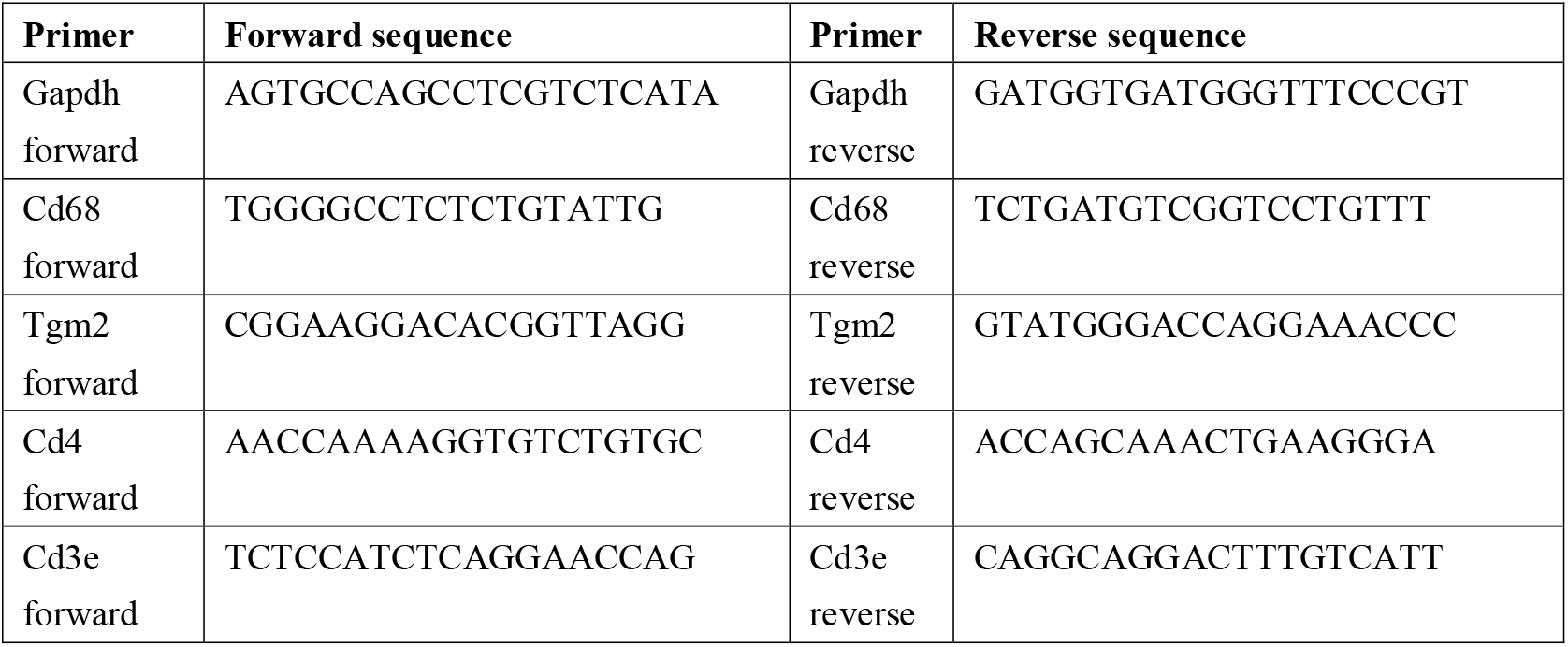
TB Green rat mRNA primers.

### Fluorescence activated Cell Sorting (FACS) analysis

The surface markers of macrophages and their phenotypes were examined by flow cytometry to evaluate proportion and polarization of macrophages.

The in vivo specimens were first cut into pieces (diameter < 1mm). The tissue was mixed with 10ml enzyme mix containing Type I Collagenase (3125u/mL) (Gibco) and 2.5ml trypsin (Gibco), and poured into a gentleMACS™ C Tube. After dissociating the tissue on gentleMACS Dissociator for 37s (Skin mode), another 10ml enzyme mix was added. The sample was digested for 2.5-3 hours at 37□ in a rotary machine (Peqlab). Then sample was passed through a 70μm cell strainer (Corning), centrifuged at 300×g for 5 minutes at room temperature. Cells were gently washed twice with PBS containing 0.05% BSA and stored in an ice box. Then the cell solutions were co-incubated with antibodies against iNOS (PE, NBP2-22119PE, Novus), CD68 (FITC, ab134351, Abcam), Arg1 (PE/Cyanine7, Cat# 369707, Biolegend), Ly6c (Alexa Fluor 700, Cat# 128023, Biolegend) and Mrc1 (Alexa Fluor 647, ab195192, Abcam) at 1:400 dilution in the dark for 1 h at 4°C (100μl per antibody for each sample). All samples were centrifuged at 450RCF for 5 min at 4°C. Supernatants were removed by aspiration, 1ml 1XPBS solution containing 0.05% BSA was used to wash the cells twice. FACS analysis was performed on NovoCyte Flow Cytometers (ACEA Biosciences®, San Diego, California) and FlowJo10.5.0. The experiments were repeated for three times independently (n=3).

### Histological and immunofluorescent staining

The sections were pretreated with 1% BSA in PBS containing 0.1% Triton X 100 for 1 h, incubated in 1% Tween 20 for 20 min and washed again in PBS. The sections were subsequently analyzed for Krt10 and Krt5, according to the manufacturers’ instructions. Briefly, sections were incubated for 30 min in dark. The excessive dye was rinsed off with PBS. Sections were incubated with antibody isotype to exclude false positive staining. Double immunofluorescence staining with primary antibodies against cytokeratin 10 (ab76318, Abcam, 1:150), cytokeratin 5 (ab52635, Abcam, 1:200) and secondary antibodies (GB25303, GB21303, Servicebio, 1:400) was performed. The immunostained specimens were further subjected to Hoechst33258 staining (G1011, Servicebio). At least three parallel sections were observed with fluorescence microscope (ZEISS SteREO Discovery.V20, Olympus). The fluorescence area measurement was conducted on five random sights of regenerated epithelium with CaseViewer 2.1 and Image Pro Plus 7.0 (n = 5).

### Statistical Analysis

Statistical significance for in vivo and in vitro data of histological analysis, qPCR and FACS were analyzed by analysis of variance (ANOVA) at the 95% confidence level, which were performed in GraphPad Prism 8.0 (GraphPad Software, San Diego, CA, USA) and P < 0.05 was considered statistically significant, while P > 0.05 was considered having no statistical differences, which was marked with NS.

